# Pyk2 in dorsal hippocampus plays a selective role in spatial memory and synaptic plasticity

**DOI:** 10.1101/2021.05.02.442340

**Authors:** Vincenzo Mastrolia, Omar al Massadi, Benoit de Pins, Jean-Antoine Girault

## Abstract

Pyk2 is a Ca^2+^-activated non-receptor tyrosine kinase enriched in the forebrain, especially in pyramidal neurons of the hippocampus. Previous reports suggested its role in hippocampal synaptic plasticity and spatial memory but with contradictory findings possibly due to experimental conditions. Here we address this issue and show that novel object location, a simple test of spatial memory induced by a single training session, is altered in Pyk2 KO mice and that re-expression of Pyk2 in the dorsal hippocampus corrects this deficit. Bilateral targeted deletion of Pyk2 in dorsal hippocampus CA1 region also alters novel object location. Long term potentiation (LTP) in CA1 is impaired in Pyk2 KO mice using a high frequency stimulation induction protocol nut not with a theta burst protocol, explaining differences between previous reports. The same selective LTP alteration is observed in mice with Pyk2 deletion in dorsal hippocampus CA1 region. Thus, our results establish the role of Pyk2 in specific aspects of spatial memory and synaptic plasticity and show the dependence of the phenotype on the type of experiments used to reveal it. In combination with other studies they provide evidence for a selective role of non-receptor tyrosine kinases in specific aspects of hippocampal neurons synaptic plasticity.

## Introduction

Synaptic function and plasticity are regulated by multiple signaling pathways including numerous protein kinases, among which evidence supports a role for non-receptor tyrosine kinases of the Src family (SFKs) and protein tyrosine kinase 2b (Ptk2b) commonly known as Pyk2. Pyk2 is a non-receptor tyrosine kinase closely related to focal adhesion kinase (FAK, Ptk2) but in contrast to FAK, Pyk2 is activated by increased cytoplasmic free Ca^2+1^. The mechanism of action of Ca^2+^ may include facilitation of Pyk2 dimerization^2, 3^. Depending on cell types Pyk2 activation requires the activity of PKC^4^, Ca^2+^/calmodulin-dependent protein kinases^5^ or calcineurin^6^, but whether these enzymes act directly or indirectly is not known. Pyk2 is highly expressed in the forebrain, especially in hippocampus^7^ where it is activated by neuronal activity^2, 8-10^.

Pyk2 and SFKs are functionally linked because activated Pyk2 autophosphorylates at Thr402, and recruits and activates SFKs (review in^11^). SFKs potentiate the activity of Pyk2 by phosphorylating its activation loop^12^. SFKs and presumably Pyk2 itself can phosphorylate and modulate NMDA receptors (NMDAR, review in^13^). Pyk2 can also interact with many protein partners including PSD-95, which binds Pyk2 proline-rich motif through its Src-homology domain 3 (SH3)^14^. This may anchor Pyk2 to NMDAR with which PSD-95 interacts through its PDZ domain (review in^15^). Interestingly Ca^2+^ can induce the dimerization of PSD-95 and trigger the autophosphorylation of Pyk2^2^. Glutamate and NMDA cause post-synaptic clustering of Pyk2, which is prevented by overexpression of PSD-95 SH3 domain^2^. The upregulation of Grin2A (NMDAR subunit 2A, a.k.a. NR2A) induced by SFKs involves PSD-95 phosphorylation and its interaction with Pyk2^16^.

Several reports provided evidence for a role of Pyk2 in synaptic plasticity, a process in which tyrosine phosphorylation has been known for many years to be implicated^17^. Pyk2 mediates the upregulation of NMDAR currents induced by metabotropic glutamate receptor 1 (Grm1, a.k.a. mGluR1) in cortical neurons in culture^18^. In cultured hippocampal neurons brain-derived neurotrophic factor (BDNF) increases NMDAR currents by enhancing local translation of Pyk2^19^. Pyk2 potentiates NMDAR currents in acutely dissociated hippocampal CA1 pyramidal neurons, whereas overexpression of kinase-dead Pyk2 impairs long term potentiation (LTP) in this preparation^10^. Interfering with Pyk2 interaction with PSD-95 also prevents LTP induction^2^. In contrast, in cultured hippocampal slices, Pyk2 knock-down blocks long term depression (LTD) but not LTP induction^20^. These results support a role of Pyk2 in synaptic plasticity but the differences suggest that this role may be dependent on experimental conditions. Recent studies investigated synaptic plasticity in Pyk2 KO mice with diverging conclusions. We reported that in the absence of Pyk2 LTP is altered at Schaffer collaterals-CA1 pyramidal cells synapses^21^, whereas another group using a different Pyk2 KO mouse line found normal LTP but impaired LTD at these synapses^22^. It is important to underline that in these various studies, the protocols used for studying synaptic plasticity were not the same (see Results and Discussion). In addition, the two studies in Pyk2 KO mice reported different results for hippocampal-dependent memory with impairment of novel object location^21^, but no alteration in the Morris water maze^22^.

In order to clarify the functional role of Pyk2 and the differences in findings between various reports we examined novel object location (NOL), a simple and sensitive spatial memory test^23^, in mice with a constitutive Pyk2 KO or a selective Pyk2 deletion in the dorsal hippocampus. We also re-examined synaptic plasticity at CA1 synapses and compared in the same mouse line the protocols previously used by different groups in different lines. We show that mice with a constitutive Pyk2 deletion display a NOL deficit that is rescued by Pyk2 re-expression and mimicked by targeted deletion in the dorsal hippocampus. We also show that the LTP alteration in slices from mice with a Pyk2 KO or its targeted deletion in CA1 neurons depends on the protocol used for inducing potentiation. Our observations indicate that Pyk2-deficient mice display a specific memory alteration and clarify the role of Pyk2 in synaptic plasticity showing that it is partial and dependent on the experimental protocol used for its induction.

## Results

### Spatial memory is altered in Pyk2 knockout mice

We first used a constitutive Pyk2 KO (*Ptk2b*^*-/-*^) mouse line and tested their behavioral phenotype using the novel object location test (NOL, **Figure 1A**), a task that assesses hippocampus-dependent spatial memory by testing the mouse ability to recognize the repositioning of a familiar object^24, 25^. We chose this test, which involves a single brief session of training (10 minutes) and is sensitive to dorsal hippocampus alterations^23^, because it was previously reported to be altered in Pyk2 KO mice^21^. In contrast, other commonly used tests, such as novel object recognition (NOR) and Morris water maze, which involve several brain regions or repeated training, respectively^23^ do not appear to be altered in Pyk2 KO mice^2, 26^. In the NOL paradigm, as expected, 24 hours after the first presentation, wild type animals spent more time exploring the object moved to the new location than the other object that remained at the old location. In contrast, after 24 hours Pyk2 KO animals did not show any preference for the objects at either location (**Figure 1B**). The mean difference between NL WT and NL KO was -10.9 [95.0 % CI -17.3, -4.47] with a P value of the two-sided permutation t-test of 0.004 (**Supplementary Table 1**). These results confirmed the existence of a spatial memory deficit in Pyk2 KO mice as evaluated by the NOL test.

**Figure 1:**
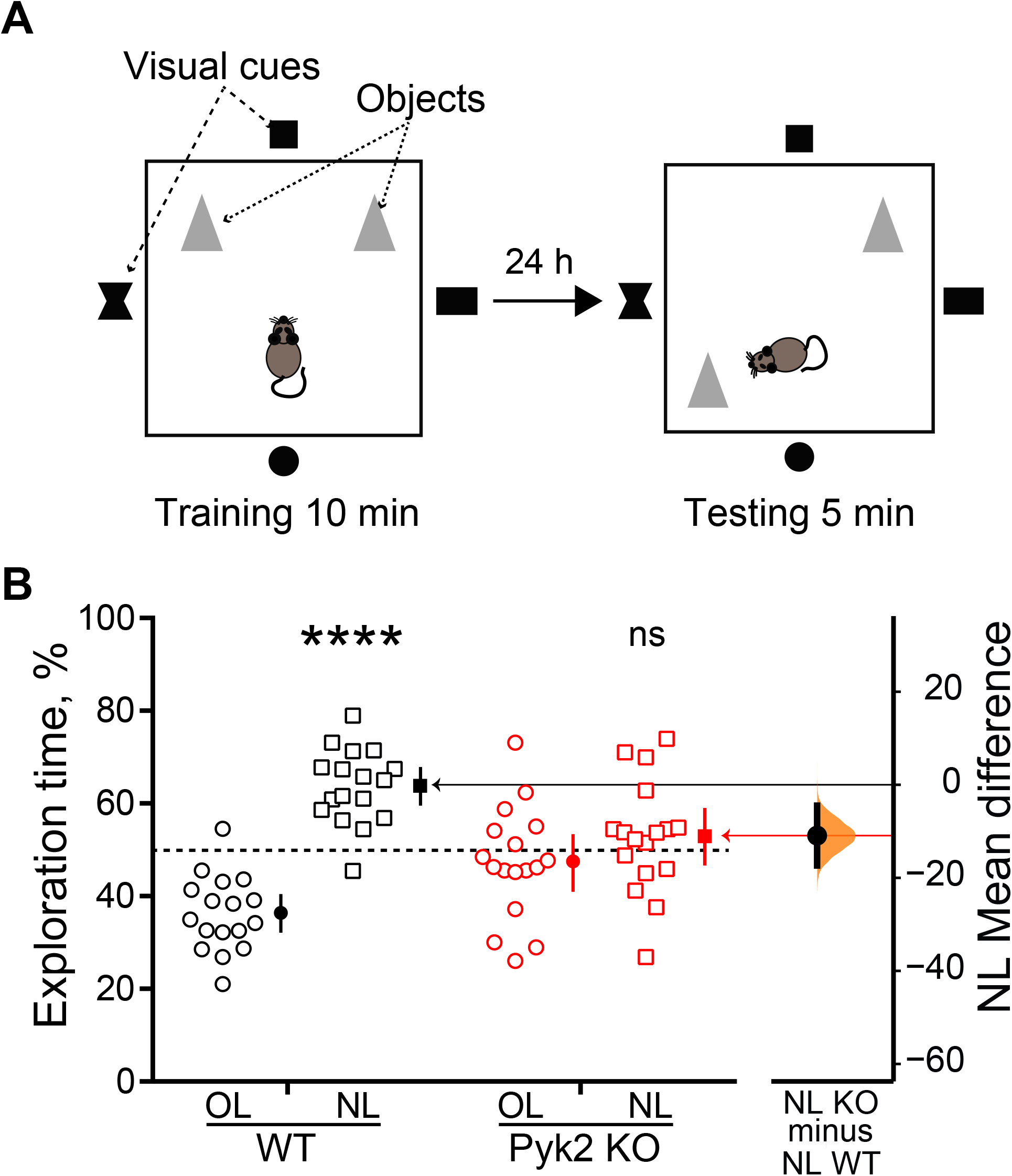
Pyk2 knockout alters novel object location (NOL) **A**)Schematic representation of the test principle. After habituation of the mouse to the arena, two identical objects (indicated as grey triangles) are positioned and the animal is allowed to explore them for 10 min (left panel). Black on white visual cues outside of the arena facilitate spatial location of the objects. One day later the mouse is placed for 5 min in the same arena in which the location of one of the two objects has been modified. The time spent exploring the displaced object (new location, NL) and the unmoved object (old location, OL) is measured by video tracking. **B**) Time spent exploring OL and NL is plotted on the left axis with open symbols for individual mouse values of Pyk2 wild type mice (n = 17) and Pyk2 KO mice (n = 17). Closed symbols indicate means, ends of vertical bars indicate the 95 % confidence interval (CI), and the horizontal dotted line indicates the chance level (50 %). One-sample t-test (NL exploration time vs. 50 % chance preference) WT, t_16_ = 6.99, p < 10^−4^, KO, t_16_ = 0.96, p = 0.35. The mean difference between NL WT and NL KO is shown as Gardner-Altman estimation plot and is plotted on a floating axis on the right as a bootstrap sampling distribution. The mean difference is depicted as a dot; the 95 % CI is indicated by the ends of the vertical error bar. Comparison of NL WT vs NL KO, unpaired two-tailed t test with Welch’s correction, t_28_ = 3.09, p = 0.0045. **, p<0.01, **** p < 10^−4^, ns, not significant. Statistical results in **Supplementary Table 1**.

### Re-expression of Pyk2 in dorsal hippocampus rescues performance of Pyk2 KO mice in the NOL test

We then thought to determine whether the deficit in the NOL test observed in the Pyk2 KO mice could be reversed by re-expression of Pyk2 in the dorsal hippocampus. To our knowledge, the reversibility of this deficit has not been investigated before. Male 2-4 month-old Pyk2 KO mice received in the dorsal hippocampus a bilateral double stereotactic injection of AAV-GFP/Pyk2 or of AAV-GFP as a control. Three weeks later they were trained for NOL. Twenty-four hours after training, the Pyk2 KO mice injected with AAV-GFP spent the same time exploring the object at the novel and old locations (**Figure 2A**), as in the case of non-injected KO mice (see **Figure 1B**). In contrast the Pyk2 KO mice injected with AAV-GFP/Pyk2 spent more time exploring the object placed at a novel location than expected by chance (**Figure 2A**). The mean time spent at NL difference between AAV-GFP and AAV-GFP/Pyk2 was 13.8 % (95.0 % CI 3.2, 24.8) with a P value of the two-sided permutation t-test of 0.024 (**Supplementary Table 1**). This result shows that the behavioral alteration observed in Pyk2 KO mice is reversible upon re-expression of Pyk2 in the dorsal hippocampus, ruling out permanent alteration or indirect effects. Because the AAV1 serotype used [^27^ and the CaM-kinase 2α (*Camk2a* gene) promoter driving the expression of Pyk2^28^ preferentially target pyramidal neurons, these results suggest that the consequences of the absence Pyk2 on NOL are linked to its deficit in dorsal hippocampus pyramidal neurons.

**Figure 2:**
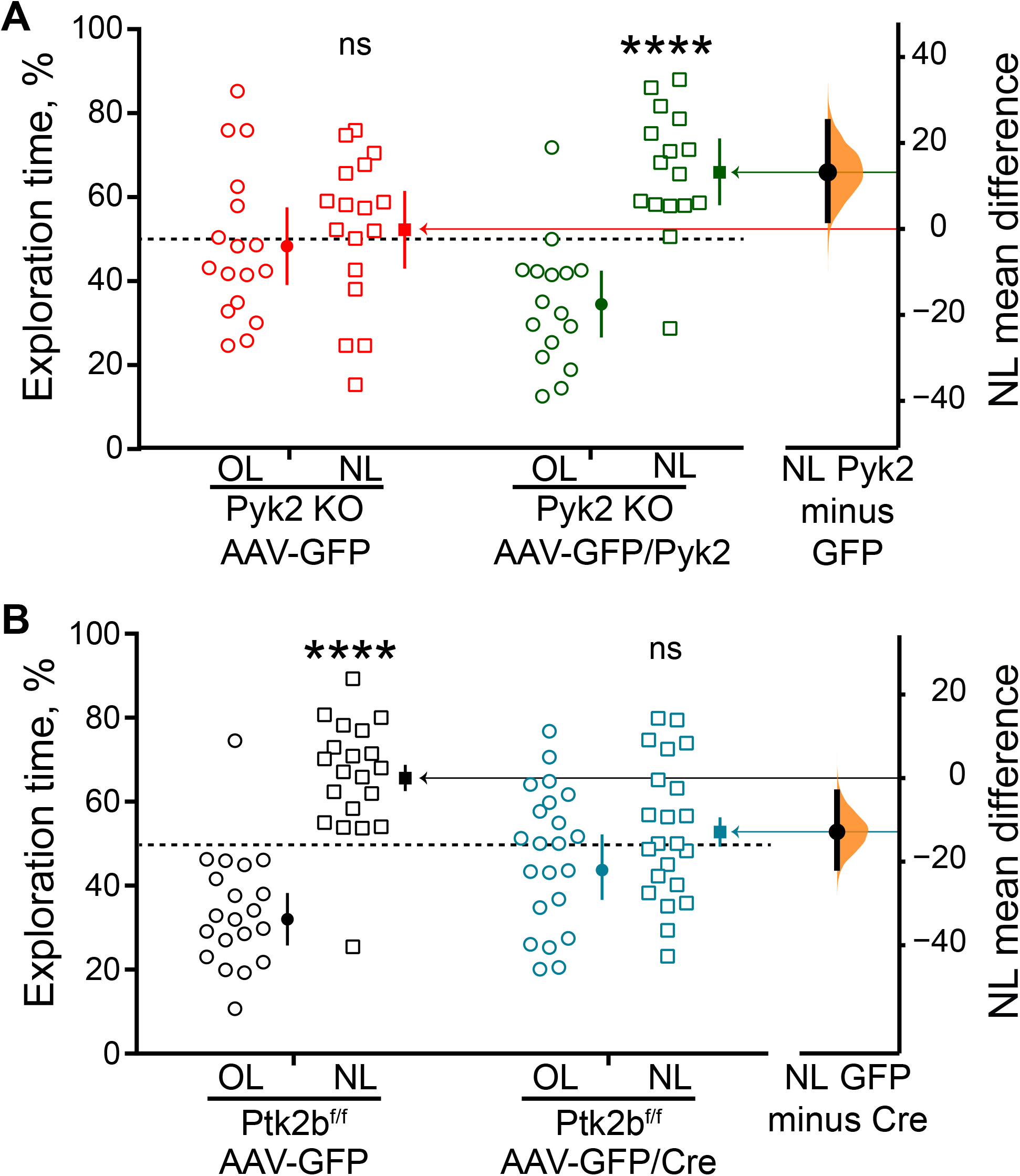
The deficit in NOL in Pyk2 KO mice is rescued by re-expression of Pyk2 in the dorsal hippocampus and mimicked by its selective deletion in this region. **A**) The deficit of Pyk2 KO mice in the NOL test is rescued by Pyk2 re-expression. Three-to-five month-old male Pyk2 KO mice were injected bilaterally in dorsal hippocampus CA1 with either AAV-GFP or AAV-GFP/Pyk2 and 3 weeks later their performance in the NOL test was evaluated as in Fig. 1. Time spent exploring OL and NL is plotted on the left axis with open symbols for individual mouse values of Pyk2 KO mice injected with AAV-GFP (n = 17) or with AAV-GFP/Pyk2 (n = 16). The time spent exploring the object at NL for AAV-GFP-injected mice was not different from random choice for KO AAV-GFP-injected mice (one-sample t-test NL exploration time vs. 50 % chance preference, t_16_ = 0.45, p = 0.66), whereas it was for KO AAV-GFP/Pyk2-injected mice (one-sample t-test t_15_ = 4.22, p = 0.0008. Comparison of NL KO AAV-GFP vs KO AAV-GFP/Pyk2, unpaired two-tailed t test with Welch’s correction, t_30_ = 2.40, p = 0.02. The mean difference between the time spent exploring NL for AAV-GFP- and KO AAV-GFP/Pyk2-injected KO mice is plotted on the right as a bootstrap sampling distribution (mean difference depicted as a dot, 95 % confidence interval indicated by the ends of the vertical error bar, two-sided permutation t-test, p = 0.024). **B**) Targeted conditional deletion of Pyk2 in dorsal hippocampus CA1 alters performance of mice in the NOL test. Three-to-five month-old male Pyk2^f/f^ were injected bilaterally with either AAV-GFP or AAV-GFP/Cre and tested for NOL three weeks later. Time spent exploring OL and NL is plotted on the left axis with open symbols for individual values for Pyk2^f/f^ mice injected with AAV-GFP (n = 20) or with AAV-GFP/Cre (n = 22). One-sample t-test (NL exploration time vs. 50 % chance preference) AAV-GFP-injected mice, t_19_ = 5.11, p < 10^−4^, AAV-GFP/Cre-injected mice, t_21_ = 0.84, p = 0.41. The mean difference between the time spent exploring NL for AAV-GFP- and AAV-GFP/Pyk2-injected mice is plotted on the right as a bootstrap sampling distribution (mean difference depicted as a dot, 95 % confidence interval indicated by the ends of the vertical error bar, two-sided permutation t-test, p = 0.0092). In A) and B), the horizontal dotted line indicates the chance level (50 %). In **A**) and **B**), bars indicate means + SEM, *, p<0.05, **** p < 10^−4^, ns, not significant. Statistical results in **Supplementary Table 1**.

### Targeted deletion of Pyk2 in dorsal hippocampus neurons of adult mice alters spatial memory

We then examined whether the alteration in NOL observed in Pyk2 KO mice could be mimicked by the targeted deletion of Pyk2 in the dorsal hippocampus. We used *Ptk2b*^*f/f*^ mice and stereotactically injected adeno-associated virus (AAV) expressing either the recombinase Cre fused to GFP (AAV-GFP/Cre) or GFP alone (AAV-GFP) in the dorsal hippocampus at two injection sites. The AAV serotype (AAV1) and the human synapsin-1 (*SYN1* gene) promoter controlling Cre expression preferentially target pyramidal neurons^27, 29^. Three weeks after the injection, we assessed spatial memory with the NOL test. Twenty-four hours after the first exposure to the objects, *Ptk2b*^*f/f*^ mice injected with AAV-GFP retained their ability to distinguish the moved object (**Figure 2B**). In contrast, the mice injected with AAV-GFP/Cre lost this ability (**Figure 2B**). The mean difference in time spent exploring NL between AAV-GFP-and AAV-GFP/Cre-injected mice was -12.9 % [95.0 % CI -21.5, -3.41] with a P value of the two-sided permutation t-test of 0.0092 (**Supplementary Table 1**). This result, in agreement with a previous report^21^, shows that targeted deletion of Pyk2 in the dorsal hippocampus is sufficient to alter performance in the NOL test. This result rules out developmental effects or indirect consequences of Pyk2 deletion in other brain regions or tissues. In addition, because the AAV serotype and the promoter of the viral vector used display cell type specificity, these results also provide information about the location of Pyk2 critical for the behavioral deficit, presumably dorsal hippocampus pyramidal neurons.

### CA1 LTP is impaired in Pyk2 KO mice in an induction protocol-dependent manner

We then explored in mice used for behavioral evaluation LTP at synapses between Schaffer collaterals and CA1 pyramidal cells, location of one of the best characterized forms of NMDA-dependent synaptic plasticity important for spatial memory^30^. We used acute hippocampal slices from wild type (age range 2.1-4.6 months, mean ± SEM, 3.0 ± 0.3) and Pyk2 KO (age range 2.0-3.2 months, mean ± SEM, 2.5 ± 0.2 months) mice. The absence of Pyk2 protein was verified by immunoblotting of the hippocampus of KO mice as compared to WT (**Figure 3A**). We stimulated Schaffer collaterals and measured fEPSP slopes in CA1. To ensure that the loss of Pyk2 had no influence on the efficiency of stimulation on the Schaffer collaterals or major pre-synaptic effect, we first analyzed the input/output relationship between the fiber volley amplitude (representing the stimulation of the Schaffer collaterals) and the fEPSP slope (representing the post-synaptic stimulation) and found no difference between wild type and Pyk2 KO slices (**Figure 3B**).

**Figure 3:**
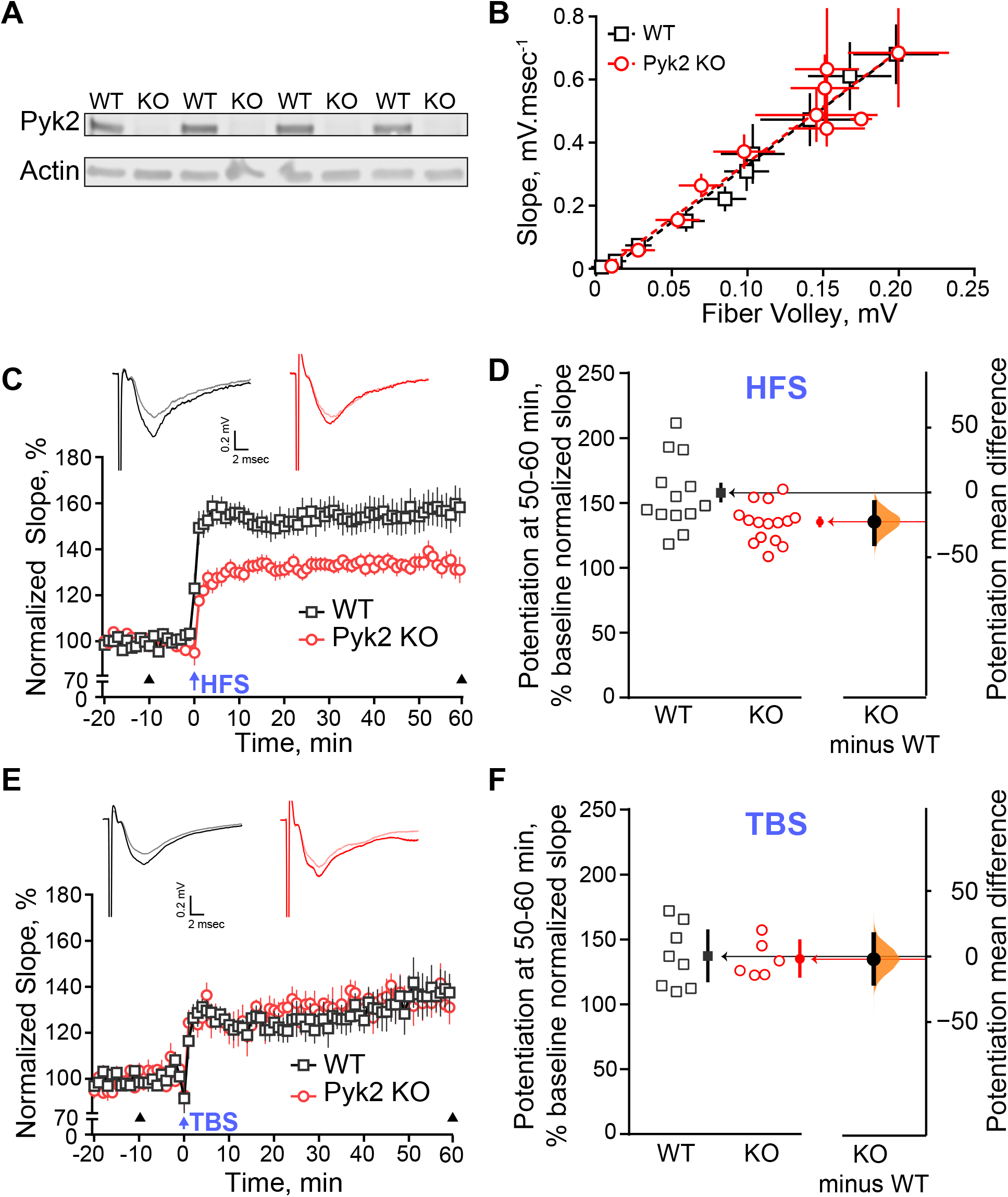
Selective impairment of LTP at CA1 synapses in Pyk2 KO mice depends on the induction protocol. fEPSP were recorded in CA1 of acute dorsal hippocampus slices from wild type (WT) and Pyk2 KO mice. Schaffer collaterals were stimulated. **A**) Hippocampi contralateral to that used for electrophysiology were obtained from 3 KO mice and 3 WT mice and analyzed by immunoblotting for the presence of Pyk2 and actin as a loading control. **B**) Input/output relationship in WT (n = 13) and Pyk2 KO (n = 15) slices (means ± SEM). No difference was observed. **C**) High frequency stimulation (HFS)-induced LTP is altered in Pyk2 KO mice. Slices from WT (8 animals, 13 slices) and Pyk2 KO (7 animals, 15 slices) mice were subjected to HFS (five 1-s trains of 100 Hz stimulation separated by 10 s intervals). Slopes of evoked fEPSPs were recorded every minute before and after HFS (arrow, time 0) and expressed as a percentage of slope average during the 20-min baseline (data for each time point are means ± SEM). Insets at the top: representative traces 10 min before and 60 min after HFS stimulation (arrowheads). Two-way repeated measure ANOVA, interaction, F_(79, 2054)_ = 4.63, p < 10^−4^, genotype effect, F_(1, 26)_ = 15.76, p = 5 10^−4^. **D)** Potentiation during the last 10 min of recording in (**C**). The average normalized slope expressed as % of baseline was calculated for each slice 50-60 min after HFS induction and plotted as open symbols with the closed symbols and the vertical bars indicating the group mean and its 95 % confidence interval (**left axis**). The mean difference between the WT and Pyk2 KO slices is plotted on the **right axis** as a bootstrap sampling distribution (mean difference depicted as a dot, 95 % confidence interval indicated by the ends of the vertical error bar, two-sided permutation t-test, p = 0.0092). **E**) Theta burst stimulation (TBS)-induced LTP is not altered in Pyk2 KO mice. Hippocampal slices from WT (5 animals, 8 slices) and Pyk2 KO (5 animals, 6 slices) mice were stimulated using a TBS protocol (fifteen 4-pulse bursts at 100 Hz, every 200 msec). Time course of normalized slopes of fEPSPs were recorded and plotted as in (**C**), before and after TBS (time 0). Insets at the top: representative traces 10 min before and 60 min after TBS stimulation (arrowheads). Two-way repeated measure ANOVA, interaction, F_(79, 948)_ = 0.63, p > 0.99, genotype effect, F_(1, 12)_ = 0.10, p = 0.75. **F**) Potentiation at the end of the recording in (**E**), expressed as in (**D**). Two-sided permutation t-test, p = 0.855. In **D**) and **E**), * p < 0.05, *** p < 0.001. **C-F**, statistical results in **Supplementary Table 1**.

In these experimental conditions, we induced LTP using two different stimulating protocols, which have been previously used to assess the role of Pyk2 in hippocampal synaptic plasticity with contrasting results^21, 22^. The first protocol was a high-frequency stimulation (HFS) protocol [5 times 1 s stimulation at 100 Hz with an interval of 10 s^21^], and the second protocol was a theta-burst stimulation (TBS) protocol [10 bursts of 4 shocks at 100 Hz with an interburst interval of 200 ms^22^]. When we induced LTP in slices from wild type animals with the HFS protocol we elicited a robust and persistent LTP (**Figure 3C** and **D**). In contrast, when we stimulated slices from Pyk2 KO animals with HFS, we found a significantly reduced induction of short term potentiation (0-10 min after HFS and LTP (**Figure 3C** and **D, Supplementary Table 1**). The mean difference in average EPSP slope at 50-60 min (expressed as % of baseline normalized slope) between Pyk2 WT and KO mice was -22.6 % [95.0 % CI -39.6, -7.65], with a two-sided permutation t-test p value = 0.0068 (**Figure 3D, Supplementary Table 1**). During the same experimental sessions and using different slices, including some from the same animals as for the HFS protocol, we induced LTP using the TBS protocol described above. With TBS we found no difference in the elicited LTP between slices from wild type or Pyk2 KO animals (**Figure 3E** and **F, Supplementary Table 1**). The mean difference in average EPSP slope at 50-60 min between Pyk2 WT and KO mice was -2.1 % (95.0 % CI, -20.5, 16.8), with a two-sided permutation t-test p value = 0.86 (**Figure 3F, Supplementary Table 1**). These results obtained with two different types of stimulation in the same mouse line show that the alteration in LTP induced by the absence of Pyk2 depends on the protocol used for inducing synaptic potentiation.

### Targeted deletion of Pyk2 in dorsal hippocampus CA1 alters LTP in an induction protocol-dependent manner

We then examined whether loss of Pyk2 limited to dorsal hippocampus CA1 region was able to affect synaptic plasticity in acute hippocampal brain slices, as above for the NOL test. We measured fEPSPs in CA1 of hippocampal slices from *Ptk2b*^*f/f*^ animals injected with either AAV-GFP/Cre (age range 2.4-6.4 months, mean ± SEM, 4.9 ± 0.6) or AAV-GFP (age range 2.4-6.6 months, mean ± SEM, 4.6 ± 0.8) and measured the LTP induced by either HFS or TBS. We confirmed the correct targeting of the stereotaxic injections by in each slice on the recording rig (**Figure 4A, left panel** and **a1-2**) and observed that GFP fluorescence was confined to CA1, mostly in the pyramidal layer (**Figure 4A, a3-5**). We analyzed the input/output relationship between fiber volley amplitude and fEPSP slope in slices from *Ptk2b*^*f/f*^ mice injected either with AAV-GFP/Cre or AAV-GFP and found no difference (**Figure 4B**), supporting the lack of presynaptic effect, as expected by the restriction of viral spread to CA1.

**Figure 4:**
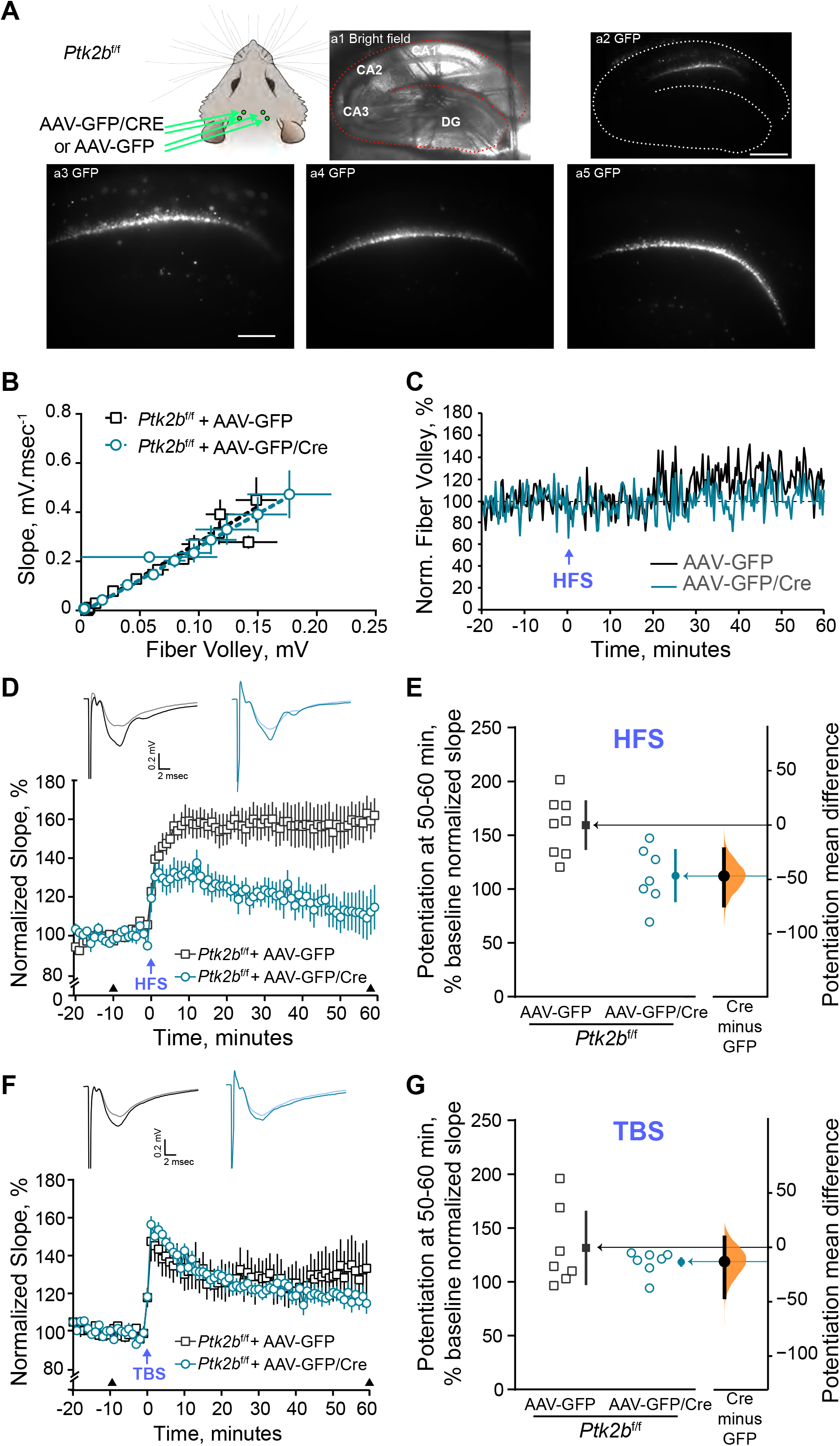
Targeted conditional deletion of Pyk2 in the dorsal hippocampus results in a selective induction-dependent impairment of LTP at CA1 synapses. **A**) *Ptk2b*^*f/f*^ mice were injected bilaterally at 2 sites in dorsal hippocampus CA1 region with either AAV-GFP/Cre or AAV-GFP (**top left panel**). Successful expression was evaluated with fluorescence microscopy on slices prior to electrophysiological recordings (**a1**, bright field, **a2**, GFP fluorescence, **a3** same as a2 at higher magnification (a1-3 are from the same slice), **a4** and **a5**, different mice). Scale bar, a1-2, 500 µm, a3-5 200 µm. (**B-E**) fEPSP were recorded as in Fig. 3, in CA1 of acute hippocampal slices from Pyk2^f/f^ mice injected with AAV-GFP or AAV-Cre/GFP with stimulation of Schaffer collaterals. **B**) Input/output relationship in slices from Pyk2^f/f^ mice injected with AAV-GFP (n = 8) or AAV-GFP/Cre (n = 7, means ± SEM). No difference was observed. **C**) Time course of fiber volley amplitude (normalized to the mean before HFS) during the whole recording in slices stimulated with HFS. **D**) High frequency stimulation (HFS)-induced LTP is altered in mice with a Pyk2 deletion in dorsal hippocampus. Slices from Pyk2^f/f^ mice injected with AAV-GFP (4 animals, 8 slices) or AAV-GFP/Cre (4 animals, 7 slices) were subjected to HFS as in Fig. 3B. Insets at the top: representative traces 10 min before and 60 min after HFS stimulation (arrowheads). Two-way repeated measure ANOVA, interaction, F_(79, 1027)_ = 5.13, p < 10^−4^, Cre effect, F_(1, 13)_ = 19.83, p = 0.004. **E**) Potentiation during the last 10 min of recording in (**D**). The average normalized slope expressed as % of baseline was calculated for each slice 50-60 min after HFS induction and plotted as open symbols with the closed symbols and the vertical bars indicating the group mean and its 95 % confidence interval (**left axis**). The mean difference between the WT and Pyk2 KO slices is plotted on the **right axis** as a bootstrap sampling distribution (mean difference depicted as a dot, 95 % confidence interval indicated by the ends of the vertical error bar, two-sided permutation t-test, p = 0.0064). **F**) Theta burst stimulation (TBS)-induced LTP is not altered by selective deletion of Pyk2 in the dorsal hippocampus. Hippocampal slices from Pyk2^f/f^ mice injected with AAV-GFP (5 animals, 7 slices) or AAV-GFP/Cre (4 animals, 7 slices) mice were stimulated using a TBS protocol as in Fig. 3C. Time course of normalized slopes of fEPSPs were recorded as in (**D**) before and after TBS (time 0). Insets at the top: representative traces 10 min before and 60 min after TBS stimulation (arrowheads). Two-way repeated measure ANOVA, interaction, F_(79, 948)_ = 0.89, p = 0.73, Cre effect, F_(1, 12)_ = 0.30, p = 0.59. **G**) Potentiation during the last 10 min of recording in (**F**), expressed as in (**E**). Two-sided permutation t-test, p = 0.41. **D-G**, statistical results in **Supplementary Table 1**.

We then induced LTP in different slices with either HFS or TBS, the two protocols described above. We first monitored the fiber volley amplitude over time after HFS and did not observe any major change or difference between AAV-GFP- and AAV-GFP/Cre-injected mice (**Figure 4C**), supporting the absence of presynaptic modification. As expected, in slices from *Ptk2b*^*f/f*^ animals injected with AAV-GFP, the HFS protocol induced a robust and persistent LTP (**Figure 4D**) at levels comparable to wild type animals (**Figure 3C**), showing the lack of effect of the presence of floxed alleles of *Ptk2b* gene, virus injection or GFP expression on this parameter. In contrast, LTP induced by HFS was markedly reduced in slices from *Ptk2b*^*f/f*^ mice injected with AAV-GFP/Cre (**Figure 4D, Supplementary Table 1**). The mean difference in average EPSP slope at 50-60 min (expressed as % of baseline normalized slope) between slices of GFP- and GFP/Cre-expressing mice was -46.9 % [95.0 % CI -74.0, -22.2], with a two-sided permutation t-test p value = 0.0064 (**Figure 4E, Supplementary Table 1**). We also examined the effects of the TBS protocol in slices from Pyk2^f/f^ animals injected with AAV-GFP or AAV-GFP/Cre. TBS elicited similar levels of LTP in slices from *Ptk2b*^*f/f*^ animals injected with AAV-GFP/Cre or AAV-GFP (**Figure 4F, Supplementary Table 1**). The mean difference in average EPSP slope at 50-60 min between slices of GFP/Cre- and GFP-expressing mice was -13.1 % [95.0 % CI - 45.9, 9.07], with a two-sided permutation t-test p value = 0.41 (**Figure 4G, Supplementary Table 1**). These results showed that targeted deletion of Pyk2 in dorsal hippocampus CA1 neurons replicates the effects of constitutive knockout, with a selective deficit in LTP induced with a HFS protocol and no alteration with the TBS protocol. They show that the HFS-induced LTP modification is not the consequence of a developmental or indirect alteration. Moreover because the AAV was expressed in CA1 pyramidal layer neurons and not in CA3 or other cell types, these results indicate that the alteration in HFS-induced LTP very probably results from a post-synaptic defect in these neurons.

## Discussion

In this study, we provide evidence that the loss of the non-receptor tyrosine kinase Pyk2 in mice alters spatial memory and synaptic plasticity in hippocampal neurons in a selective manner. These results support previous observations indicating a role of Pyk2 in spatial memory and LTP and provide a rationale for explaining the phenotypic differences between reports in different mouse lines, which displayed either deficits in NOL and LTP^21^ or normal LTP and spatial memory in the Morris water maze^22^. We confirm the alteration in the NOL test as reported by Giralt et al (2017) but not explored by Salazar et al. (2019). This deficit was rescued by re-expression of Pyk2 in the dorsal hippocampus CA1 region and reproduced by targeted deletion in neurons of this brain area. These findings rule out that the phenotypic deficit results from developmental alterations or indirect effects. In addition, because the viral vector serotype (AAV1)^27^ and the promoters used to drive the expression of Pyk2 (mouse *Camk2a* gene)^28^ and Cre (human *SYN1* gene)^29^ are known to preferentially target hippocampal pyramidal neurons, this memory deficit is likely to derive from the absence of Pyk2 in pyramidal neurons, where it is abundantly expressed^7^. It is important to underline that the memory deficits in Pyk2 KO mice are only partial with altered NOL [^21^ and this report], but normal performance in the Morris water maze^22^. The Morris water maze is a classical test with multiple variations that explores spatial memory and is altered by hippocampal lesions or dysfunction^31, 3^. The NOL test, which evaluates spatial memory 24 h after a single exposure, is highly sensitive to and relatively selective for dorsal hippocampal alterations [see^23^ for a recent review]. We do not know why Pyk2-deficient mice display a deficit in the NOL but not in the water maze test, but it is worth noting that the NOL involves a single training session whereas the water maze protocol used with Pyk2 KO mice included multiple training sessions during several days and may be reinforced by the stressful component of the test^33^. We suggest that the partial impairment of synaptic plasticity in Pyk2 mutant mice is sufficient to alter one-trial NOL performance but can be overcome by repeated training or possibly a stress-linked component. Interestingly, a different one-trial training test, normal novel object recognition (NOR), was reported to be normal in Pyk2 KO mice^21, 2^. Although NOL and NOR may appear similar their sensitivity to interferences is different^23^. NOR involves several brain regions and neurotransmitter systems^34^. Aggleton and Nelson conclude that “transient disruptions of dorsal hippocampal activity and plasticity are sufficient to impair NOL, though the same manipulations spare NOR”^23^. The selective deficit in Pyk2 is likely to be compensated by other mechanisms, including involvement of other brain regions, when memory is tested in NOR but not in NOL. These results highlight that various commonly used behavioral tests explore different aspects of the memory processes and that a selective molecular perturbation can alter some of them but not others.

The selectivity of Pyk2 requirement in hippocampal function is also highlighted by its implication in synaptic plasticity. Several reports using entirely *in vitro* approaches provided indications of a role of Pyk2 in LTP at hippocampal synapses. Huang and colleagues carried out patch-clamp studies in acutely dissociated CA1 pyramidal neurons and found that Pyk2 in the patch pipette potentiated NMDA currents^10^. The presence of kinase-dead Pyk2 in the pipette prevented the LTP induced by a tetanic stimulation consisting of two 500-ms trains of 100 Hz stimuli at a 10-sec interval^10^. Bartos and colleagues also investigated LTP in acute hippocampal slices using whole cell patch clamp and tetanic stimulation with two 500-ms 100 Hz trains separated by 10 s^2^. They reported that LTP induction was prevented when the patch pipette contained the SH3 domain of PSD-95 that binds Pyk2 (GST–SH3) or the Pyk2 sequence with which this SH3 domain interacts (GST–Pyk2[671-875]). The authors concluded that Pyk2 interaction with PSD-95 was necessary for the induction of LTP in these conditions, presumably allowing Pyk2 synaptic clustering and activation^2^. A third report investigated synaptic plasticity in hippocampal slices in organotypic culture with whole cell patch clamp^20^. The authors used a different type of protocol to induce LTP by pairing a stimulation consisting in 200 pulses at 3 Hz with postsynaptic depolarization to 0 mV and found no effect of Pyk2 down-regulation with shRNA^20^. Divergent results were also reported in studies with mutant mice. LTP induced in hippocampal slices from Pyk2 KO mice at Schaffer collaterals synapses on CA1 pyramidal cells by 5 times repeated 1-s tetanus at 100 Hz with an interval of 10 s was blocked^21^. In contrast, in a different Pyk2 KO mouse line, LTP induced at the same synapses by a theta-burst stimulation (TBS) protocol (10 bursts of 4 shocks at 100 Hz with an interburst interval of 200 ms) was not altered^22^. Here we compared these two protocols of LTP induction in the same Pyk2 KO mouse line, including in different slices from the same mice and found that the LTP induced by HFS was altered whereas LTP induced by TBS was not modified. We carried out the same experiments in *Ptk2b*^*f/f*^ mice injected with AAV-GFP/Cre in the dorsal hippocampus and observed that HFS-induced LTP was impaired whereas TBS-induced LTP was uchanged in comparison to AAV-GFP-injected mice. Thus, in two different mouse models we replicated the previous findings of the two different groups, providing evidence that the difference in their results was linked to the use of different LTP induction protocols. Both direct manipulation of Pyk2 in hippocampal neurons^2, 10^ or complete or targeted Pyk2 KO (Giralt et al., 2017 and present report) alter HFS-induced LTP, whereas LTP induced by TBS (Salazar et al., 2019 and present report) or by a related stimulation protocol^20^ is not altered. Interestingly it was reported that HFS-induced LTP was also impaired in heterozygous Pyk2-KO mice (Giralt et al., 2017), suggesting a dosage sensitivity. In the present work we did not further explore this point since we used homozygous constitutive Pyk2 KO mice and in the virally-induced deletion it was not possible to know whether the deletion was homozygous in all targeted cells.

The importance of the stimulation protocol is also apparent in the retrospective comparison of other studies on the role of tyrosine kinases in synaptic plasticity. The inhibitory effects of tyrosine kinase inhibitors on LTP at CA1 synapses were observed with HFS^17, 35^. Grant et al. investigated the hippocampal LTP at CA1 synapses in four non-receptor tyrosine kinase KO lines, 3 SFKs (Fyn, Src, and Yes), and the structurally more distant Abl^36^. They observed a deficit in Fyn KO mice but not in the other mouse lines, and only when they used two trains of tetanic stimulation trains (1-s trains at 100 Hz) with an intertrain interval of 20 s^36^. In contrast they did not observe any alteration when LTP was induced by pairing 40 low frequency (1 Hz) stimulations with post-synaptic depolarization^36^. This protocol is reminiscent of the one that did not reveal any LTP alteration in Pyk2 knockdown slices (200 pulses at 3 Hz paired with postsynaptic depolarization), as mentioned above^20^. Taken together these results indicate that the role of non-receptor tyrosine kinases Pyk2 and Fyn at synapses between Schaffer collaterals and CA1 pyramidal neurons is limiting for specific types of plasticity and can be overcome in other conditions.

Why is the role of Pyk2 important for HFS-induced LTP and not TBS-induced LTP? Previous work from many laboratories provided evidence that HFS-induced LTP results from the pronounced increase in Ca^2+^-influx through NMDAR [review in^30^]. TBS, also implicates NMDAR activation, but its late phase specifically requires the activation of cAMP-dependent protein kinase^37-39^, illustrating the differential contribution of various signaling pathways to different aspects of synaptic plasticity. Several reports show that Pyk2 can modulate the intensity of NMDAR currents^10^ through the recruitment of SFKs [see^13^]. The converging evidence showing the importance of SFKs and Pyk2 in HFS-induced LTP indicates that these tyrosine kinases are specifically required for this type of synaptic plasticity. The activation of Pyk2 and its links with SFKs including Fyn is depicted in **Figure 5**. Because tyrosine phosphorylation decreases endocytosis of NMDAR [review in^40^], one hypothesis to explain the role of Pyk2 and SFKs is that they oppose this endocytosis during intense activation and thereby maintain sufficient Ca^2+^ influx. Our data show the consequences of the absence of Pyk2 which could be a consequence of the absence of its tyrosine kinase activity, or possibly of some other properties, such as protein-protein interaction. It will be interesting to explore these aspects by using acute and specific pharmacological inhibition of Pyk2 tyrosine kinase activity.

**Figure 5:**
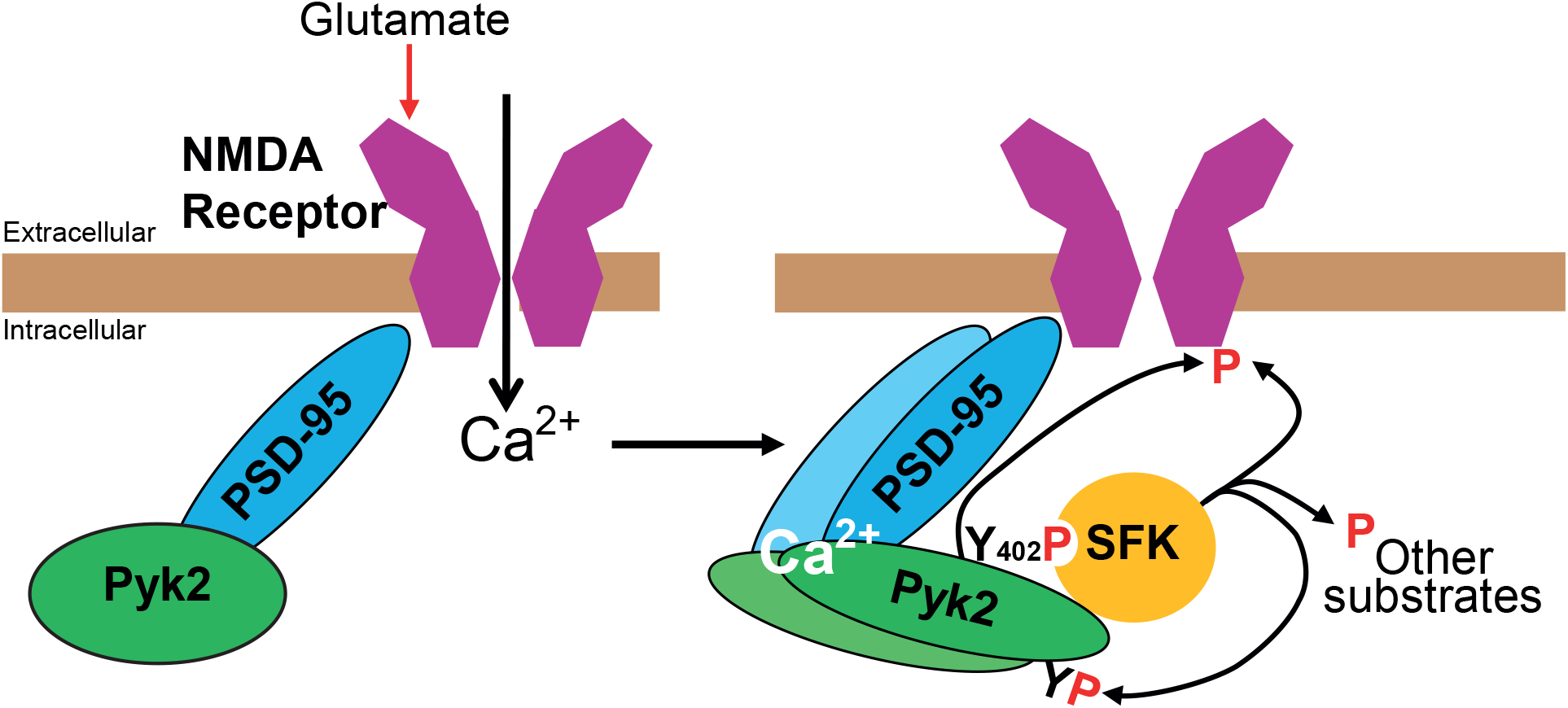
Schematic representation of Pyk2 activation at synapses. Pyk2 is associated with NMDA receptors (NMDAR) through its interactions with PSD protein PSD-95. Following Ca^2+^ entry through activated NMDAR, Pyk2 is activated, presumably by intermolecular autophosphorylation promoted by interaction of Ca^2+^ with Pyk2 and/or with associated protein such as PSD-95. Autophosphorylated Tyr-402 in Pyk2 provides a binding site for the Src-homology domain 2 (SH2) of Src-family kinases (SFKs) such as Fyn or Src. SFKS also interact with Pyk2 through their SH3 domain (details not depicted in the cartoon). This results in the activation of SFKs which can phosphorylate various tyrosine residues in Pyk2 and increase its kinase activity. SFKs and presumably Pyk2 phosphorylate NMDAR on tyrosine, which increases NMDA currents, at least in part by increasing NMDAR at the cell surface. In addition, SFKs and Pyk2 can phosphorylate other substrates and activate various signaling pathways that have not yet been fully characterized at the post-synaptic level. See the text for references.

Importantly Pyk2 also plays a role in LTD. LTD in CA1 pyramidal neurons induced by pairing a 200-pulse stimulation at 1 Hz with depolarization of the postsynaptic cell to -40 mV is markedly decreased by Pyk2 knockdown^20^ and LTD induced in CA1 by a 900-pulse stimulation at 1 Hz of Schaffer collaterals is prevented in Pyk2-KO mice^22^. The pathways mediating these two forms of LTD are therefore altered in the absence of Pyk2. Interestingly tyrosine phosphorylation of GluA2 subunit of AMPA receptors was shown to be important for LTD^41^. However, since there are several overlapping types of LTD coexisting at these synapses involving mGluR1 activation and decreased tyrosine phosphorylation as well as NMDAR-dependent endocytosis of AMPA receptors [review in^42^], further work is needed to elucidate the precise molecular mechanisms of Pyk2 contribution to LTD.

In conclusion, the present work clarifies the role of Pyk2 in synaptic plasticity and memory. It confirms that in the absence of Pyk2 in the dorsal hippocampus a one-trial test of spatial memory, NOL, is impaired and shows that this impairment can be rescued by re-expression of Pyk2. The lack of alteration of other tests^21, 2^ indicates that the consequences of Pyk2 deficit can be overcome by repeated training (Morris water maze) or recruitment of extra-hippocampal mechanisms (novel object recognition). Our results also demonstrate the dissociation between the role of Pyk2 in LTP, which is necessary for HFS-induced LTP but not TBS-induced LTP in CA1. This dissociation is in agreement with previous reports on the role of non-receptor tyrosine kinases at those synapses. Our findings clarify previous contradictions between results of several laboratories and provide a better understanding of the role of non-receptor tyrosine kinases in synaptic plasticity. Future work is needed to elucidate the precise molecular mechanisms underlying the selective requirement of Pyk2 for specific aspects of memory and synaptic plasticity.

## Material and Methods

### Mice

All experiments were carried out in male mice. We used *Ptk2b*^*-/-*^ (Pyk2 KO) mice and mice with floxed Pyk2 alleles (*Ptk2b*^*f/f*^) on a C57Bl/6 background. KO mice were bred by mating *Ptk2b*^*+/-*^ heterozygous mice. The *Ptk2b*^*-/-*^ (KO) and *Ptk2b*^*+/+*^ (wild type) used for experiments originated from these breeding pairs. Mice were housed under conditions of controlled temperature (23°C) and illumination (12-hour light/12-hour dark cycle, ZT0 = 7:00 am). All animal procedures were authorized by and performed in accordance with the animal care committee’s regulations.

All experiments were carried out in male mice. We used Ptk2b-/-(Pyk2 KO) mice and mice with floxed Pyk2 alleles (*Ptk2b*^*f/f*^) on a C57Bl/6 background^21, 43^. KO mice were bred by mating *Ptk2b*^*+/-*^ heterozygous mice. The Ptk2b^-/-^ (KO) and *Ptk2b*^*+/+*^ (wild type) used for experiments originated from these breeding pairs. Mice were housed under conditions of controlled temperature (23°C) and illumination (12-hour light/12-hour dark cycle, ZT_0_ = 7:00 am). All animal procedures were conducted in accordance with guidelines of Declaration of Helsinki and NIH (1985-revised publication no. 85-23, European Community Guidelines), and French Agriculture and Forestry Ministry guidelines for handling animals (decree 87849, license A 75-05-22), with approval of the Charles Darwin ethical committee (agreement APAFIS#8861-2016111620082809).

### Viral constructs and stereotaxic injection

For specific deletion of Pyk2 in the dorsal hippocampus, 6-10 week-old male Pyk2^f/f^ mice were stereotactically injected either with AAV expressing eGFP fused with the recombinase Cre, including a Woodchuck hepatitis virus posttranscriptional regulatory element (WPRE) to enhance expression, under the control of the human synapsin-1 gene promoter (AAV-GFP/Cre, 105540-AAV1, pENN.AAV1.hSyn.HI.eGFP-Cre.WPRE.SV40, Addgene, Massachusetts, USA) or with AAV expressing only GFP (AAV-GFP, AV-9-PV1917, AAV9.CamKII0.4.eGFP.WPRE.rBG from Perelman) as control. For re-expression of Pyk2 in the dorsal hippocampus of Pyk2 KO mice, the AAV used was AAV1-CamKIIα (0.4)-GFP-2AmPTK2B, polycistronically expressing GFP and Pyk2 separated by a 2A site (AAV-GFP/Pyk2, Vector Biolabs Malvern, PA, USA) or, as control, AAV-GFP, as above. Mice were anesthetized by an intraperitoneal injection of ketamine 15 mg kg^-1^ and xylazine 3 mg kg^-1^, and placed in a stereotaxic frame (Kopf Instruments, Sweden). The hippocampus was targeted bilaterally using a 32-gauge needle connected to a 1-ml syringe (Neuro-Syringe, Hamilton). Two injections were done on each side, at different locations in the dorsal hippocampus CA1 region with the following coordinates from the bregma (millimeters): AL -1.8, ML ±1.2, DV -1.5, and AL -2.3, ML ±1.5, DV -1.5. AAVs were delivered at a rate of 0.1 μl min^-1^ for 5 min (0.5 μl per injection site). After the viral particles were infused, the needle was kept in place for 5 additional minutes before removal. After the procedure, the incision was closed with surgical sutures, and mice were placed in a heated cage until they recovered from anesthesia. After 2 h of monitoring, mice were returned to their home cage for 3 weeks before starting analyses of behavior and electrophysiology. To reduce suffering, mice received a subcutaneous injection of a non-steroidal anti-inflammatory drug (flunixin, 4 mg/kg) just after the surgery and on the next day, when necessary.

### Behavioral phenotyping

For novel object location (NOL) test we used an open-top arena (45 × 45 x 45 cm) under dim light (60-70-lux intensity). Mice were first habituated to the arena for 2 days for 15 min per day. To allow mice to identify their position, 4 different visual cues were placed around the arena: one circle, one square, one rectangle and one with the shape of an hourglass, all solid black on a white background. On day 3, two identical objects (Eiffel tower bronze figurines, painted in gray, about 9 cm-high and 3 cm in diameter) were placed in the arena and mice were left to explore the arena for 10 min. Twenty-four hours later, one of the objects was moved from its original location to the diagonally opposite corner and mice were allowed to explore the arena for 5 min (**Figure 1A**). The mice were continuously recorded with a video tracker (Viewpoint, Newcastle Upon Tyne, UK) placed above the arena, and the percentage of time exploring the displaced object (new location, NL) and the unmoved object (old location, OL) was compared. Object exploration was defined as mice actively sniffing or touching the object with their nose or forepaws. Quantification was done manually with a stopwatch.

### Slice preparation

Male 2-6 month-old mice were anaesthetized by an intraperitoneal injection of ketamine 15 mg kg^-1^ and xylazine 3 mg kg^-1^. Following the achievement of full anesthesia and analgesia, mice where immobilized and a sagittal thoracic incision was made in order to expose the heart. An ice-cold low Ca^2+^/high Mg^2+^ artificial cerebrospinal fluid (aCSF) solution (pH 7.4, 310 mOsm, in mM, 92 NaCl, 26 NaHCO_3_, 1 NaH_2_PO_4_, 3 KCl, 12.5 glucose, 20 Hepes, 0.2 CaCl_2_, 7 MgCl_2_,) was intracardially perfused for 2-3 minutes until the liver acquired a pale color. The mouse was then swiftly decapitated, the brain was removed and quickly placed in a custom-made 3D-printed brain matrix to rapidly excise the hemispheres with a 45°-angle, necessary for producing hippocampal transversal slices^44^. Each hemisphere was placed on a Petri dish, hold in place with magnets on the vibrating slicer cutting block (Thermo Fisher, USA) and filled with ice-cold low Ca^2+^/high Mg^2+^ aCSF solution with constant carbogen (95 % O_2_/5 % CO_2_) bubbling. Slices (350 μm-thick) were cut (0.08 mm s^-1^, blade vibration, 60 Hz and 1 mm) and individually moved in NMDG-aCSF solution (pH 7.4, 310 mOsm, in mM, 93 N-Methyl-D-glucamine [glucamine, NMDG], 30 NaHCO_3_, 1.2 NaH_2_PO_4_, 2.5 KCl, 25 glucose, 20 Hepes, 7 ascorbate, 4 pyruvate, 0.2 CaCl_2_, 7 MgCl_2_ [adapted from^45^] for 10 min at 32° C for recovery. Slices were then moved to a custom-made 3D-printed interface chamber^44^ containing a modified aCSF solution (pH 7.4, 310 mOsm, in mM, 92 NaCl, 26 NaHCO_3_, 1 NaH_2_PO_4_, 3 KCl, 12.5 glucose, 20 Hepes, 2 CaCl_2_, 1.3 MgCl_2_) with constant carbogen (95 % O_2_/5 % CO_2_) bubbling and allowed to rest for 2 hours at room temperature.

### Electrophysiology

After the 2-hour rest, each slice was placed in a custom-made 3D-printed perfusion chamber^44^, with constant flow of recording artificial CSF solution (aCSF, pH 7.4, 310 mOsm, in mM, 124 NaCl, 26 NaHCO_3_, 1 NaH_2_PO_4_, 3 KCl, 12.5 glucose, 5 Hepes, 2.5 CaCl_2_, 1 MgCl_2_) at 32° C with constant carbogen (95 % O_2_/5 % CO_2_) bubbling and let equilibrate for 30 minutes. The stimulating and recording electrodes were made of silver wire encased in borosilicate glass pulled pipettes (pipette resistance: stimulating electrode 1-1.5 MΩ, recording electrode 5-6 MΩ) filled with aCSF. The electrodes were positioned in the hippocampal CA1 area, in the middle of the *stratum radiatum*, 300 μm apart from each other. Stimulating currents were delivered using and external ISO-Flexi stimulator (A.M.P.I., Israel) while field excitatory post-synaptic potential (fEPSP) acquisition was performed with a 700B amplifier (Molecular Devices, USA) connected to a 1440a digitizer (Molecular Devices, USA) using the Clampex 10 acquisition software (Molecular Devices, USA). To stimulate the Schaffer collaterals coming from the CA3 area, the stimulating electrode (positioned proximal, towards CA3) delivered a single depolarizing current pulse. A recording electrode was placed distal (towards the subiculum) within the *stratum radiatum* to record the fEPSPs elicited by Schaffer collaterals stimulation. The current amplitude was set to 50 % of the current that elicited the maximal fEPSP amplitude (current injection 0.6-1.2 μA). Stimulating pulses (0.1 ms) were delivered every 20 seconds (0.033 Hz) and the resulting fEPSPs were recorded. After collecting a 20-minute baseline, LTP was induced with either a high-frequency stimulation (HFS) protocol consisting of 5 tetanic bursts of 1 sec at 100 Hz separated by 10-second intervals or with a theta-burst stimulation (TBS) consisting of 15 bursts of 4 spikes at 100 Hz separated by 200-millisecond intervals. After the induction, fEPSPs were monitored during the next 60 minutes and LTP was quantified by analyzing the slope of the activation phase. Only slices with stable recordings throughout the experiment and, in the case of those injected with AAV, only slices with good GFP expression in CA1, were included in the final analyses.

### Immunoblotting

Tissue samples were sonicated in 10 g/L SDS and placed at 100⍰°C for 5⍰min. Extracts (15⍰μg protein) were separated by SDS–PAGE and transferred to nitrocellulose membranes (GE Healthcare, Chicago, IL, USA). Membranes were blocked in TBS (150⍰mM NaCl, 20⍰mM Tris-HCl, pH 7.5) with 0.5⍰ml⍰L^−1^ Tween 20 TBS-T) with 30⍰g L^−1^ BSA. Immunoblots were probed with rabbit polyclonal antibodies for Pyk2 (#P3902, Sigma-Aldrich) diluted 1:1000. Membranes were incubated with the primary antibody overnight at 4⍰°C by shaking in TBS. After several washes in TBS-T, blots were incubated with secondary anti-rabbit IgG DyLight™ 680 conjugated antibodies (1:10,000; Rockland Immunochemicals, Pottstown, PA, USA). Secondary antibody binding was detected by Odyssey infrared imaging apparatus (Li-Cor Inc., Lincoln, NE, USA).

### Statistical analyses

Data were analyzed with d’Agostino-Pearson and Shapiro-Wilk normality tests. When their distribution did not significantly deviate from normal distribution and the number of replicate was sufficient (>7) we used Student’s t test for comparing two groups. If these conditions were not satisfied we used non-parametric Mann-Whitney’s test. Comparison of 4 groups were done with 2-way ANOVA and time courses were analyzed with 2-way repeated measures ANOVA, as indicated. Post hoc multiple comparisons after ANOVA were done with Holm-Sidak’s test. All tests were used as two-tailed. Statistical analyses were done with GraphPad Prism 6. Estimation statistics was done as recommended^46^ using the https://www.estimationstats.com/ website^47^. Statistical analyses are reported in **Supplementary Table 1**.

## Supporting information

Supplemental Table 1, detailed statistical analyses

## Acknowledgements

The authors thank Nicolas Gervasi for his help in the initial set up of electrophysiological experiments.

## Author Contributions

JAG, VM, OaM designed research; VM, OaM, BdP performed experiments; VM, OaM, BdP, JAG analyzed data; VM, OaM, JAG wrote the paper; all authors reviewed the manuscript.

